# Raw milk for human consumption may carry antimicrobial resistance genes

**DOI:** 10.1101/853333

**Authors:** Adrienn Gréta Tóth, István Csabai, Eszter Krikó, Dóra Tőzsér, Gergely Maróti, Árpád V. Patai, László Makrai, Géza Szita, Norbert Solymosi

## Abstract

**Background:** The increasing prevalence of antimicrobial resistance (AMR) is a significant threat to global health. The widespread use of antibiotics is increasingly shortening the time it takes for resistant strains to develop. More and more multi-drug-resistant bacterial strains cause life-threatening infections and the death of tens of thousands of people each year. Beyond disease control animals are often given antibiotics for growth promotion or increased feed efficiency, which further increase the chance of the development of multi-resistant strains. After the consumption of unprocessed animal products, these strains may meet the human bacteriota. Among the foodborne and the human populations, antimicrobial resistance genes (ARGs) may be shared by horizontal gene transfer. This study aims to test the presence of antimicrobial resistance genes in milk metagenome, investigate their genetic position and their linkage to mobile genetic elements.

**Results:** We have analyzed raw milk samples from public markets sold for human consumption. The milk samples contained genetic material from various bacterial species and the detailed analysis uncovered the presence of several antimicrobial resistance genes. The samples contained complete ARGs influencing the effectiveness of acridine dye, cephalosporin, cephamycin, fluoroquinolone, penam, peptide antibiotics and tetracycline. One of the ARGs, PC1 beta-lactamase may also be a mobile element that facilitates the transfer of resistance genes to other bacteria, e.g. to the ones living in the human gut.

**Conclusion:** Besides the animal products’ antibiotic residuals, their potentially transmissible antimicrobial resistance gene content may also contribute to the development of human pathogenic bacteria’s antimicrobial resistance.

## Background

The increasing prevalence of antimicrobial resistance (AMR) is a significant threat to global health. The widespread use of antibiotics, both in human healthcare and animal disease control [1–3], is increasingly shortening the time it takes for resistant strains to develop and more and more multi-drug-resistant bacterial strains cause life-threatening infections. There is increasing evidence that antimicrobial resistance genes (ARGs) responsible for the occurrence of phenotypically expressed antimicrobial resistance are widespread in various environmental samples [4–7]. The pool of ARGs being present in a particular environmental sample is called the resistome [8]. In samples where the medical use of antibiotics can be excluded, usually, only a few ARGs are present [9, 10]. When antibiotics are extensively used for preventive and therapeutic purposes, bacterial strains respond to this selection pressure and will become increasingly resistant what finally leads to these agents’ elevated prevalence. ARGs hosted by non-pathogenic bacteria can be transferred to pathogens with horizontal gene transfer (HGT) what elevates the latter group’s resistance against antibiotics. The execution of HGT depends on several factors, albeit physical closeness of bacteria always increases the chances [11]. The likelihood of HGT is even higher if ARGs are carried on mobile genetic elements (e.g. plasmids). In order to understand the chance of an ARG’s horizontal gene transfer derived spread, studies aiming to assess the bacteria’s resistome and the specific position of the identified ARGs [12] are very well-reasoned and necessary.

The microbiota of livestock products may come to direct contact with the human bacteriota, either during the processing steps or during the consumption of these products. The antibiotics used for farm animal disease control often share active substances with human medicines. Consequently, there is a risk that ARGs accumulated as a response to the high amount of antibiotics used in livestock farming may be transmitted to the human microbiota through animal products. Such spread of ARGs may reduce the efficacy of antibiotic therapies even more and may lead to the development of new multidrug-resistant strains. Fortunately, food processing typically contains heat treatment steps that kill the majority of bacteria. Thus the role of active DNA-export mechanisms between the intestinal and the nutriment’s bacteriome is lower [13].

Raw milk is a product sold unprocessed; thus the presence or the grade of heat-treatment steps are upon the decision of consumers. Together with the preference of several other organic products, the consumption of non-heat-treated raw milk justified by its favourable health effects is commonly set as a trend in the developed societies [14, 15].

To our best knowledge, no previous study has investigated the possible presence of ARGs in raw milk, and we have found no data on the raw milk’s resistome. This study aims to test the presence of ARGs in milk metagenome, investigate their genetic position and their linkage to mobile genetic elements [16, 17].

## Results

### Metagenome

Two times one litre of raw milk was bought at a public market in Budapest (sample A) and in Szeged (sample B). After DNA extraction and sequencing (see Methods), from sample A 17,773,004 while from sample B 8,425,326 paired-end reads were recovered. By the quality filtering, 0.20% and 0.80% of the reads were discarded from sample A and B, respectively. The reads were aligned to the host (*Bos taurus*) genome. As expected, most of the genetic material originated from the milking cow, from sample A 96.41% and from sample B 97.01% of the cleaned reads were filtered out due to host origin.

Of the reads, not aligning to the cow genome, we were able to classify 42.11% in sample A and 52.96% in sample B to known taxa. 185,982 reads of sample A and 11,437 reads of sample B were identified to belong to the kingdom of Bacteria. In sample A 86.32% of the reads were classified as Gram-positive bacteria, while in sample B this proportion was only 13.68%. The detailed composition of the core bacteriomes at class level is shown in Figure 1.

**Figure 1.**
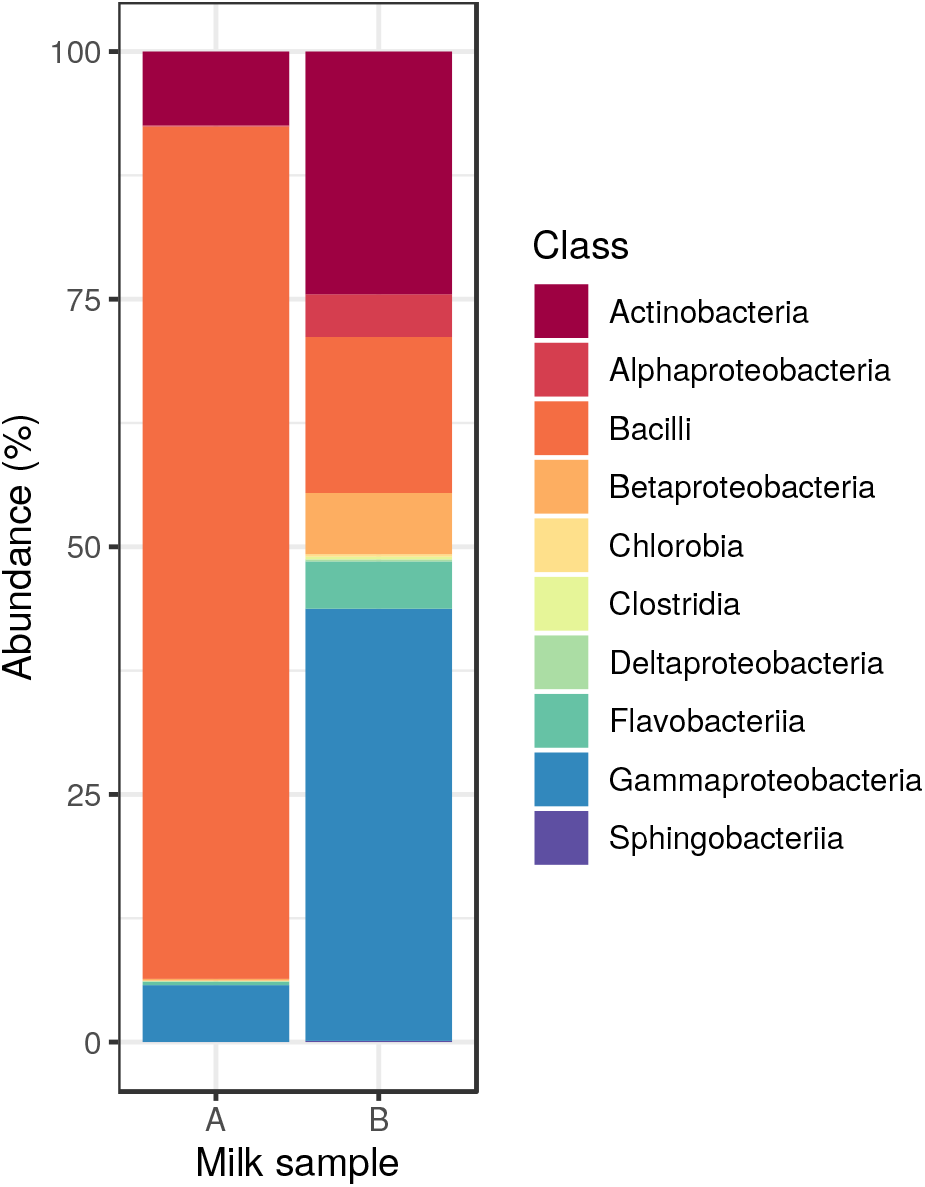
Core bacteriome composition. Relative abundance of bacteria classes by milk samples.

### ARGs and MGEs

Reads with overlapping pieces were assembled into longer DNA contigs by the metaSPAdes tool. The assembled contigs having a shorter length than 162 bp (sample A: 0.68%, sample B: 0.33% of all contigs) were excluded. The remaining contig’s median length was 268 (IQR: 126.5) and 244 (IQR: 42) in sample A and B, respectively. Then, the contigs having any open reading frame (ORF) matched with an ARG in the Comprehensive Antibiotic Resistance Database (CARD) were collected. The detected ARGs and particular properties are presented in Table 1.

**Table 1.**
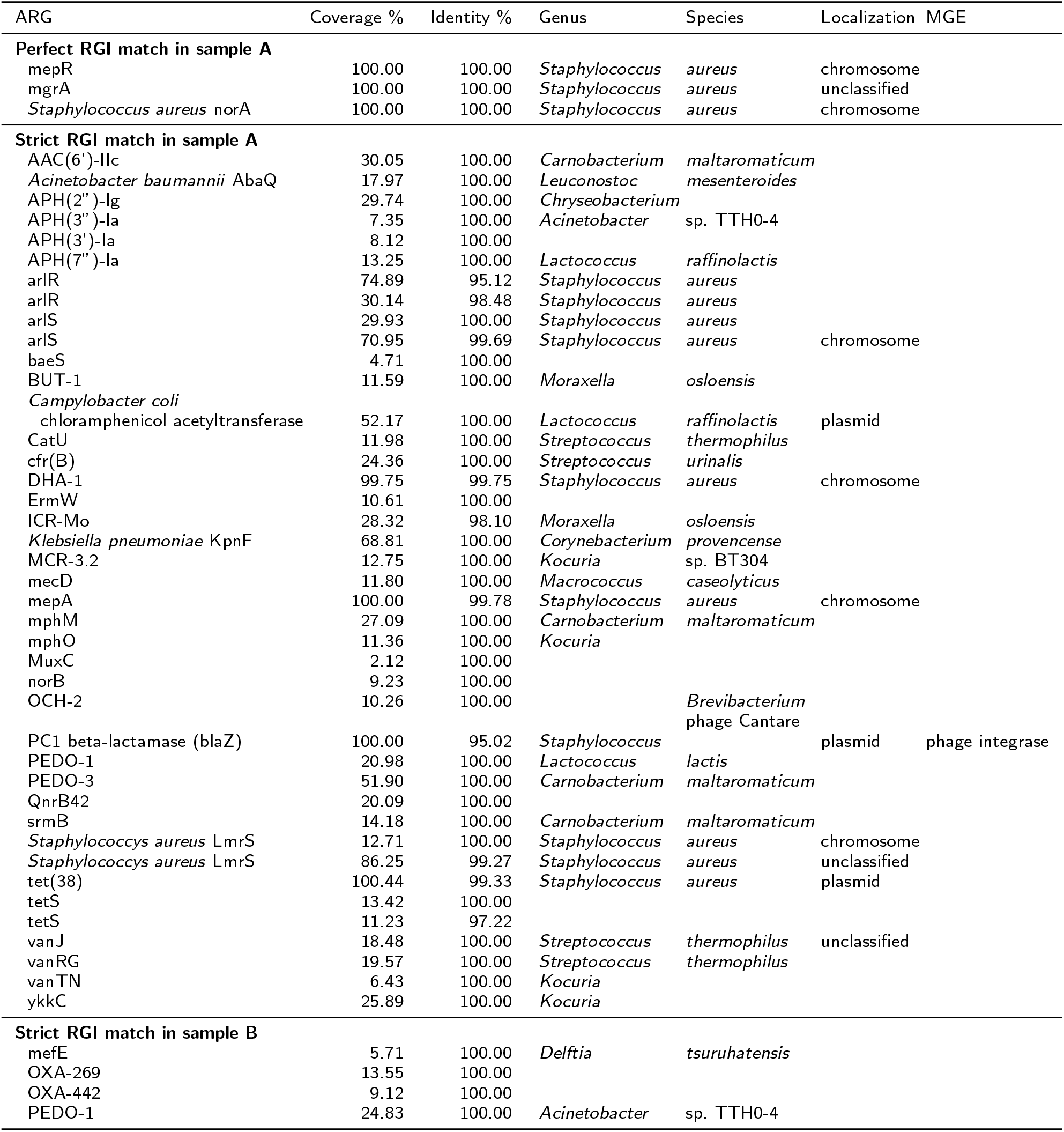
ARGs identified in milk samples. The coverage column shows the fraction of CARD ARG reference sequence covered by the most similar ORF sequence. Identity represents the proportion of the identical nucleotides between the detected ORF and CARD ARG reference sequence. Species column shows the most likely species related to the ARG harbouring contig classified by Kraken2. For some contigs, the species level classification was ambiguous, genus reported only. The localization of contigs with ARG and longer than 1000bp predicted by PlasFlow. Mobile genetic element domains coexisting with ARG are listed in column MGE.

The identified ARGs were classified with the Resistance Gene Identifier (RGI) tool according to the ratio of their coverage in the samples and to the identity between the contigs assembled from the sequenced reads and the CARD ARG reference sequences. In Table 1 we list each ARG with perfect or strict hits predicted by RGI. We were able to identify three perfect ARG matches in sample A, mepR, mgrA and *Staphylococcus aureus* norA. According to the taxonomical classification of the contigs harbouring these ORFs their most likely origin is bacteria from Staphylococcus genus. The MGE analysis showed that none of these ORFs is mobile.

The sequence coverages of the strict matches in sample A ranged between 2.12% and 100%, with a mean of 36.61%. The identity of ORFs and CARD ARG reference sequences ranged between 95.02% and 100%, with a mean of 99.59%. Contigs containing ARG were classified on genus level and Acinetobacter (2.86%), Carnobacterium (11.43%), Chryseobacterium (2.86%), Corynebacterium (2.86%), Kocuria (11.43%), Lactococcus (8.57%), Leuconostoc (2.86%), Macrococcus (2.86%), Moraxella (5.71%), Staphylococcus (37.14%) and Streptococcus (11.43%) genera were identified.

In the bacterial genome, ARGs may be located on chromosomes or on plasmids, the latter ones being more likely to translocate between bacteria. With the PlasFlow tool, we identified contigs harbouring chloramphenicol acetyltransferase, PC1 beta-lactamase (blaZ) and tet(38) ARGs that may be encoded on plasmids. The results of MGE domain coexisting analysis showed that PC1 beta-lactamase (blaZ) ARG might be mobile since the contig had a phage integrase ORF within the distance of 10 ORFs.

There were no perfect matches in sample B that is not surprising since its overall bacterial nucleic acid content was less than 10% of that of sample A. The sequence coverages of the strict matches in this sample ranged between 5.71% and 24.83%, with the mean of 13.31%. The identity between ORFs and CARD ARG reference sequences was 100.00% in each detected ARG. Contigs containing ARGs were classified and genera Acinetobacter (50%) and Delftia (50%) were identified. None of the identified ARGs could be related to any mobile genetic elements.

The detected ARGs in both samples were matched to their corresponding antibiotics. Since one antibiotic compound may be related to more than one ORFs, we decided to select those to which we could link the ORFs with the broadest coverage and the highest identity to the reference ARG sequence. The maximal coverage and identity of detected ORFs are shown in Figure 2. In sample A ARGs known to be decreasing the effectiveness of acridine dye, cephalosporin, fluoroquinolone, penam and peptide antibiotics were found in full length and with 100% identity. There were two other ARGs identified in sample A in full length and with identity above 99% that encoded resistance against further antibiotics (cephamycin and tetracycline).

**Figure 2.**
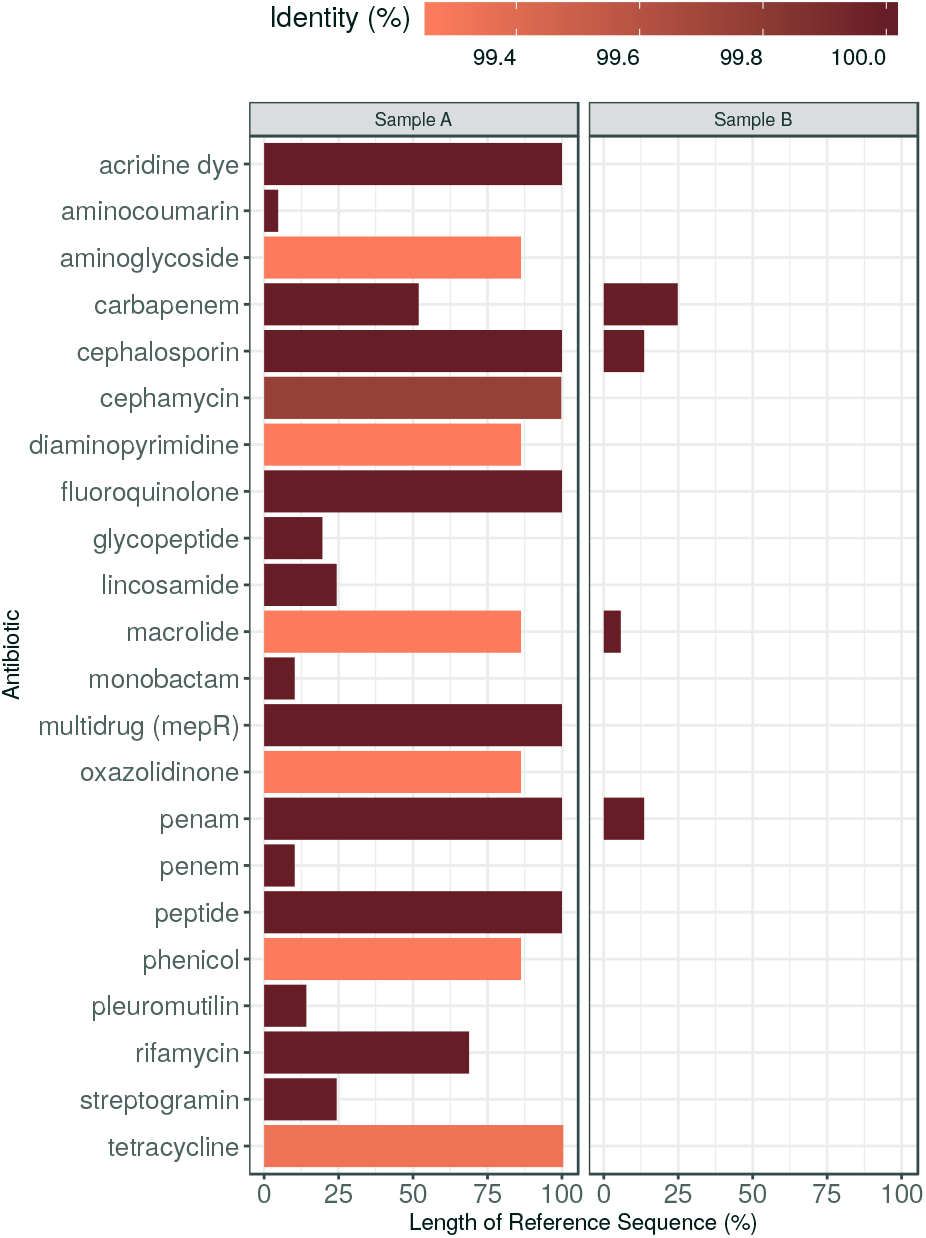
Maximal coverage and identity of detected ORFs by antibiotics. The ORF covered proportion of the reference ARG sequence (x axis) and the identity % of predicted protein (color).

## Discussion

Antimicrobial resistance (AMR) is a natural feature of microorganisms that have originally occurred as a means of defence in the rivalry amongst the members of the microbiotas. The ubiquity of antimicrobial resistance genes (ARGs) is beyond question. Genes against antibiotics are present both in non-pathogenic and pathogenic bacteria. With the extended agricultural and clinical use of antibiotics, the number of ARGs are on the rise, and the growing number and spread of multi-resistant bacteria strains pose a global threat to global health. According to the CDC’s Antibiotic Resistance Threats in the United States, 2019 report [3], more than 2.8 million antibiotic-resistant infections occur in the U.S. each year, and more than 35,000 people die as a result. In addition, 223,900 cases of *Clostridioides difficile* occurred in 2017 and at least 12,800 people died. Dedicated prevention and infection control efforts in the U.S. and around the world are working to reduce the number of infections and deaths caused by antibiotic-resistant germs. However, the number of people facing antibiotic resistance is still too high. The AR Threats Report warns that not only people but also animals carry bacteria in their guts which may include antibiotic-resistant bacteria either with intrinsic or with acquired ARGs [12]. Beyond disease control, animals may be given antibiotics for growth promotion or increased feed efficiency. Since bacteria are exposed to low doses of the drugs over a long period, this inappropriate antibiotic use can lead to the development of resistant bacteria [3]. The CDC report notes that when animals are slaughtered and processed for food, resistant germs in the animal gut can contaminate meat or other animal products, but do not mention the possible contamination of milk.

Detected ARGs in raw milk (Table 1) can be transferred from non-pathogens to pathogens via HGT. The over-expression of such genes, e.g. norA (regulated by mgrA) and mepA (regulated by mepR) coding multidrug efflux pumps confer resistance to fluoroquinolones (including norfloxacin or ciprofloxacin) and even disinfectants [18–21]. Ciprofloxacin is a broadspectrum antibiotic used to treat a variety of bacterial infections, including intra-abdominal, respiratory tract, skin, urinary tract, and bone and joint infections. Norfloxacin might be used for uncomplicated urinary tract infections (including cystitis) or the prevention of spontaneous bacterial peritonitis in cirrhotic patients, among others. MepA was also shown to result in resistance to tigecycline [22], an antibiotic that was developed to tackle complicated infections caused by multiresistant bacteria such as *Staphylococcus aureus*, *Acinetobacter baumannii*, and *E. coli*.

The two samples differ both in the composition of core bacteriome and the ARG abundance. In sample A the bacteriome was dominated by Gram-positive bacteria. Furthermore, most of the contigs harbouring ARG were classified taxonomically belonging to Grampositive bacteria. In sample B, the Gram-negative bacteria governed the bacteriome. So the lower ARG abundance in sample B might come from the lower proportion of Gram-positives. Nevertheless, in sample B, not just the number of detected ARGs was lower, but the maximal coverage of the ARGs as well. One may find the reason for this phenomenon, the lower sequencing depth of sample B. The identity of these ORFs with the reference ARGs are very high so we may assume the assembled ORFs originated from ARGs. Accordingly, the possible reason of the lower coverage of ARGs may be caused by the insufficient read counts for assembly the complete ORFs.

Our results show that indeed ARGs can be present in raw milk. However, it should be the subject of further research to identify how resistant bacterial DNA gets into the milk, is it already there in the cow’s udder or does it only mixed into the milk as contamination during or after milking.

At raw milk’s environment of origin (dairy farms), the use of antimicrobial agents is widespread. Consequently, the microbiome of this product may show relatively high levels of resistance. Without heat-treatment, bacteria that are present in raw milk are not hindered from further multiplication what results in the amplification of their resistance genes either. Such a rise in the number of ARGs may increase the risk of horizontal gene transfer (HGT) events. This risk may even be higher in case of mobile ARGs (e.g. blaZ, which was detected on a plasmid and near to a phage integrase ORF).

Beyond human intervention, there are natural mechanisms that limit ARG-transfer [11]. First of all, donor and recipient populations need to be present at the same physical space and reach a specific critical density to ensure proper connectivity for a successful gene transfer event. Chances for a series of HGT events amongst two physically distant populations are relatively low except for the case when there is positive selection driven by any factors (e.g. selection by antibiotics). The second factor arises from the fact that genes encoding resistance against the same compounds may limit each other’s spread. A population owning genes against a particular antibiotic is not under selective pressure to gain any other ARGs with the same effect. As a conclusion of earlier evolutionary steps, possession of resistance determinants of the same substrate profiles are possible. However, in a population where the distribution of these genes is stable, the chances of new recruitments are lower. Tertiary, acquisition of resistance genes sets metabolic costs deriving from the transfer and integration mechanisms needed. These costs vary by each ARG, and only affordable genes are spread [11].

Nevertheless, heat-treatment of raw milk seems to be an advantageous and a more than considerable step that besides inhibiting the amplification of genes having a potential risk, makes active gene transfer mechanisms lose their significance. On the other hand, even though it reduces the number of multiplication cycles, after the lysis of cells free DNA fragments appear in the sample that may still be uptaken by newly arriving bacteria.

However, the interpretation of resistome studies is yet to be deepened. The combination of nextgeneration sequencing, metagenomic and computational methods provides valuable data on the presence of ARGs. Moreover, it makes it possible to find genes in full coverage and length, and to identify their taxonomical classes of origin and their exact sequential surroundings. Synteny with mobile genetic elements is a fact to be taken into consideration when examining the risks meant by an ARG. Thus, the combination of methods mentioned above serves as a core component of today’s necessarily expanded antimicrobial resistance research.

## Conclusion

As a means of evolutionary pressure, the use of antibiotics selects bacterial strains that have antimicrobial resistance genes. Moreover, in the production animal sector, the application of such compounds increases not only the number of antibiotic-resistant bacterial strains but also the frequency of their appearance. After the consumption of animal products, these strains may meet the human microbiota, and the circumstances may be appropriate for the horizontal gene transfer derived spread of antimicrobial resistance genes amongst these populations. This phenomenon unfolds a possible source of acquisition of human pathogens’ antimicrobial resistance other than the direct presence of antibiotic residuals in animal products.

According to our findings, the antimicrobial resistance gene content of unprocessed animal products may play a role in the development of antimicrobial resistance of human pathogens.

## Methods

DNA extraction and metagenomics library preparation Before the laboratory procedures, the milk samples were stored frozen. 120 mL of raw milk was centrifuged at 10.000 g for 10 min. Total DNA was extracted from the pellet using the ZR Fecal DNA Kit from Zymo Research. Paired-end fragment reads (2150 nucleotides) were generated using the TG NextSeq® 500/550 Mid Output Kit v2 sequencing kit with an Illumina NextSeq sequencer. Primary data analysis (base-calling) was carried out with “bcl2fastq” software (v.2.17.1.14, Illumina).

### Bioinformatic analysis

Before the laboratory procedures, the milk samples were stored frozen. 120 mL of raw milk was centrifuged at 10.000 g for 10 min. Total DNA was extracted from the pellet using the ZR Fecal DNA Kit from Zymo Research. Paired-end fragment reads (2 150 nucleotides) were generated using the TG NextSeq® 500/550 Mid Output Kit v2 sequencing kit with an Illumina NextSeq sequencer. Primary data analysis (base-calling) was carried out with “bcl2fastq” software (v.2.17.1.14, Illumina).

Quality based filtering and trimming was performed by Adapterremoval [23], using 15 as a quality threshold. Only reads longer than 50 bp were retained. *Bos taurus* genome (ARS-UCD1.2) sequences as host contaminants were filtered out by Bowtie2 [24] with *verysensitive-local* setting minimizing the false positive match level [25]. The remaining reads were taxonomically classified using Kraken2 (*k* = 35) [26] with the NCBI non-redundant nucleotide database [27]. The taxon classification data was managed in R [28] using functions of package phyloseq [29] and microbiome [30]. For further analysis, the reads assigned to Bacteria was used only [31]. Core bacteria was defined as the relative abundance of agglomerated counts at class level above 0.1% at least one of the samples.

By metaSPAdes [32] the preprocessed reads were assembled to contigs, with the automatically estimated maximal *k* = 55.

From these contigs having a shorter length than the shortest resistance gene of the Comprehensive Antibiotic Resistance Database (CARD) were discarded [33, 34]. The ARG content of filtered contigs was analyzed with Resistance Gene Identifier (RGI) v5.1.0 and CARD v.3.0.6 [33, 34].

Contigs harbouring ARG identified by RGI with perfect or strict cut-off were preserved and classified by Kraken2 on the same way as was described above.

The plasmid origin probability of the contigs was estimated by PlasFlow v.1.1 [35].

To identify possible further mobile genetic element (MGE) homologs the predicted protein sequences of contigs were scanned by HMMER [36] against data of PFAM v32 [37] and TnpPred [38]. Following Saenz et al. [31] from the hits with lower than E 10^−5^ the best was assigned to each predicted protein within the distance of 10 ORFs. The MGE domains coexisting with ARGs were categorized as phage integrase, resolvase, transposase or transposon.

## Ethics approval and consent to participate

Not applicable.

## Consent for publication

Not applicable.

## Availability of data and material

All data are publicly available and can be accessed through the PRJNA591315 from the NCBI Sequence Read Archive (SRA).

## Author’s contributions

NS had full access to all of the data in the study and takes responsibility for the integrity of the data and the accuracy of the data analysis. LM, IC and NS conceived of the study concept. GM, LM and NS performed the sample collection and procedures. AGT, DT, EK and NS participated in the statistical analysis. AGT, AVP, LM and NS participated in the drafting of the manuscript. AGT, DT, GS, IC and NS carried out the critical revision of the manuscript for important intellectual content. All authors read and approved the final manuscript.

